# Evaluating Synergy of Clinically Utilized Phage OMKO1 with Five Antibiotics from Different Classes against *Pseudomonas aeruginosa*

**DOI:** 10.64898/2026.01.13.699299

**Authors:** Emma L. Kane, Benjamin K. Chan, Dallas L. Mould, Kaitlyn E. Kortright, Jonathan Koff, Paul E. Turner

## Abstract

Decades of antibiotic overuse and misuse have placed us at the precipice of a post-antibiotic era. This rise in multidrug-resistant and pandrug-resistant bacteria poses a serious risk to public health, warranting urgent exploration of alternative treatment options for bacterial infections. One emerging method displaying promise in the field is phage-antibiotic combination therapy: the use of bacteriophages, viruses that exclusively infect bacteria, as adjuvants to antibiotics. This study investigated the synergistic effects of phage OMKO1, previously utilized in compassionate-use cases, with five antibiotics of diverse drug classes against *Pseudomonas aeruginosa*. Modified checkerboard assays were performed to test phage-antibiotic combination treatments across several antibiotic concentrations and phage multiplicities of infection. Synergy was achieved by four of the five phage-antibiotic pairings at sub-minimum inhibitory antibiotic concentrations. All combination treatments reduced the antibiotic minimum inhibitory concentration (MIC) by ≥ 2-fold and resulted in a reduction in resistant regrowth. These findings highlight the potential of phages to lower effective antibiotic concentrations and prolong their utility by slowing the rise of antimicrobial resistance.

## Introduction

The rise of antimicrobial resistance (AMR) is a pressing health concern projected to increasingly impact public health worldwide. Since the development of penicillin in 1928, the chronic overuse and misuse of chemical antibiotics has selected for antimicrobial resistant bacteria and paved the way for a post-antibiotic era, where antibiotics are no longer able to treat many bacterial infections, specifically multidrug-resistant (MDR) bacterial infections (1,2). Organizations including the World Health Organization (WHO) have emphasized the significance of this problem, predicting that the number of morbidities caused by AMR could rise from 700,000 to 10 million by 2050 without the introduction of additional antimicrobial agents (3). The WHO has generated a list of pathogens of elevated or critical concern, which includes the ESKAPE pathogens: *Enterococcus faecium, Staphylococcus aureus, Klebsiella pneumoniae, Acinetobacter baumannii, Pseudomonas aeruginosa,* and *Enterobacter spp* (3,4). These ESKAPE pathogens are largely responsible for nosocomial infections and are increasingly dangerous for immunocompromised populations and those prone to chronic infection, for example, individuals with cystic fibrosis (CF) (5).

Caused by mutations in *CFTR* (*cystic fibrosis transmembrane conductance regulator*), CF is characterized by disrupted chloride and bicarbonate ion transport across various epithelia (6). Multiple organ systems are negatively impacted by the resulting osmotic imbalance, including the digestive, reproductive, and respiratory systems (7). In the respiratory tract, the accumulation of viscous mucus makes it difficult to expectorate passing microbes, leading to decreased mucociliary clearance and increased inflammation and infection (7). Bacteria, such as *Pseudomonas aeruginosa*, persist in the airways of people with CF (pwCF) and evolve antibiotic resistance, rendering them difficult to treat with standard anti-pseudomonal antibiotics such as inhaled tobramycin (7).

Given the rapid rise of MDR, it is imperative to explore alternative treatment options (8). One area garnering increasing attention is phage therapy: the use of lytic bacteriophages, viruses that exclusively infect and kill (lyse) bacteria, to treat bacterial infections (9–11). Unlike chemical antibiotics, bacteriophages (or phages) are extremely abundant in nature, highly specific, and self-amplifying (12–14). Upon injection of the phage genomic material, the bacterial host cell machinery is recircuited to prioritize production of more phage progeny up until lysis upon which more phage are released and ready to infect neighbouring cells. Despite these advantages, it is unlikely that phage therapy will entirely replace chemical antibiotics in the near future since bacteria are able to evolve resistance against phages as they do to antibiotics (12). However, Phage-Antibiotic Synergy (PAS), employing phages as adjuvants to antibiotics, is a promising avenue in combatting the rise of MDR (15–17). This method has been successfully used to treat MDR respiratory infections in pwCF, aortic graft infections, urinary tract infections, and more (18–23). Phage OMKO1 has specifically demonstrated the ability to re-sensitize previously resistant bacteria to antibiotics in clinical isolates and clinical uses, making it an ideal candidate for synergy testing (18). While antibiotics remain the standard of care for treating bacterial infections, the rise of compassionate-use cases which utilize bacteriophages as adjuvants to antibiotics calls for a concurrent focus on studies investigating phage-antibiotic interactions, as these combinations are not always more effective than isolated treatments (24).

Phage-antibiotic combinations can be defined in three ways: additive, synergistic, and antagonistic (25). If the combined effects are equal to the sum of either agent used alone, the combination is termed additive. If the combined effects are better (inhibit more bacterial growth) than the sum of each treatment alone, the combination is termed synergistic. Lastly, if the combination is less effective than the sum of each individual effect, the combination is antagonistic. Recent studies have suggested that these dynamics may be associated with the antibiotic’s mechanism of action: if an antibiotic facilitates cell lysis, the mechanism of lytic phages, it is reasonable to predict that the combination will be synergistic (25). Whereas, if the mode of action in some way hinders the host cell machinery responsible for DNA replication or protein synthesis, the combination could be antagonistic as the phage relies on this machinery to replicate within the cell (26).

Another consideration for developing optimal phage-antibiotic combinations is whether the antibiotic is bacteriostatic (i.e. growth inhibitory) or bactericidal (i.e. death-causing), because bacteriostatic agents may hinder the activity of bactericidal ones (27,28). Since many bacteriophages rely on exponentially growing cells for productivity, they may be best paired with bactericidal antibiotics which do not inhibit phage replication (29). Furthermore, due to the recalcitrance of *P. aeruginosa* in bloodstream and respiratory tract infections, bactericidal antibiotics are generally preferred to prevent the escalation to pseudomonal sepsis (30).

Here we present the study of five bactericidal antibiotics from different classes: ciprofloxacin, aztreonam, ceftazidime, tobramycin, and colistin, paired with clinically utilized phage OMKO1 against gram-negative *Pseudomonas aeruginosa.* Because antibiotics have been reported to trigger induction of prophages embedded in host genomes, we utilized the prophage knockout strain: MPAO1pp-(31,32). We classify each pairing as synergistic, additive, or antagonistic by using a modified checkerboard synergy assay and the Bliss Independence model of synergy. This investigation suggests that antibiotic mechanism of action is not a consistent predictor of synergy between antibiotics and phage OMKO1. Each of the combinations effectively reduced the antibiotic minimum inhibitory concentration (MIC) by ≥ 2-fold, supporting the potential of bacteriophages such as OMKO1 to limit antibiotic usage more broadly. Lastly, growth curve results from the phage-antibiotic synergy assays suggest that phage-antibiotic pairings may be accompanied by prolonged suppression of resistant regrowth, regardless of their Bliss synergy score.

## Results

### Modified Checkerboard Assays Highlight Phage-Antibiotic Interactions

To assess potential interactions between phage OMKO1 and anti-pseudomonal antibiotics, bacteria were exposed to a range of phage and antibiotic concentrations through the use of phage-antibiotic checkerboard assays (**Fig. 1**) (25). As highlighted in previous literature, this model presents three killing regions on the microplate of each assay: an antibiotic-dominated-killing region, a phage-dominated-killing region, and a phage-antibiotic interacting region (25). The first region primarily occurs where the antibiotic concentration is high enough to inhibit bacterial growth alone. The phage-dominated region occurs where the antibiotic concentration is ineffective alone, and bacterial inhibition is driven by phage. To study the interaction between phage and antibiotics, we focused on the third region of the microplate assays, the phage-antibiotic interacting region, where antibiotic and phage are present at sub-minimum inhibitory concentrations and thus largely ineffective in isolation. The wells in the phage-antibiotic interacting region were compared to control lanes which contained bacteria with antibiotic (column 11) or phage (row H) in isolation (**Fig. 1**).

**Figure 1.**
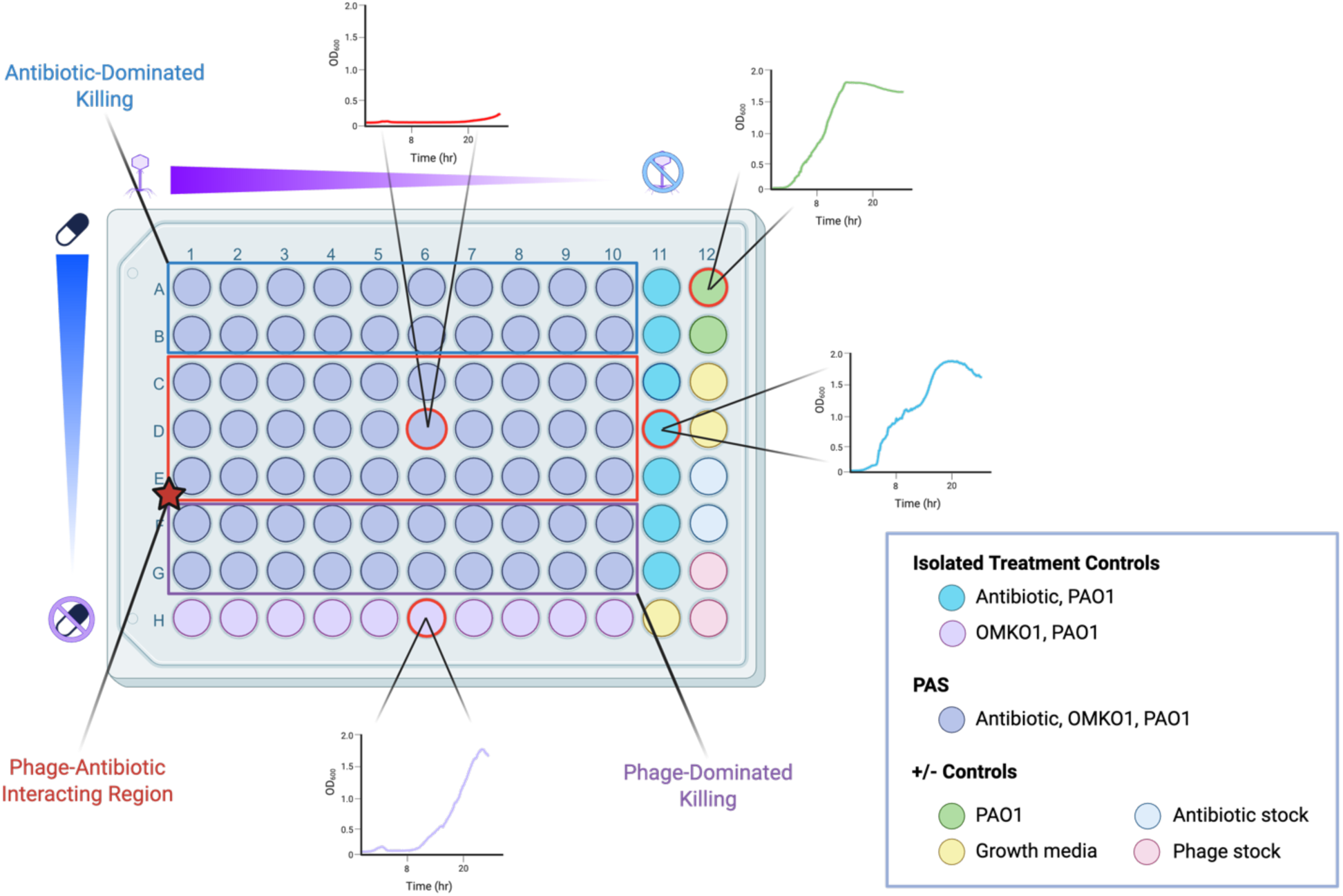
Phage-Antibiotic Synergy (PAS) assays were designed to evaluate the concentration-dependent interactions between phage and bacteria over time. In all PAS assays, row A contained the highest concentration of antibiotic, which was serially diluted 2-fold vertically down each plate. In a similar fashion, column 1 contained the highest concentration of phage and was serially diluted 1:10 horizontally along each plate. To initiate growth, a 100-fold dilution of P. aeruginosa overnight culture was added into each well, as indicated. Growth was monitored by OD_600_ readings every 10 minutes for 24 h at 37°C with constant agitation. The mean total growth of each well was captured by calculating the area under the curve over the 24-h period for three biological replicates relative to the untreated growth controls (column 12, green). To facilitate direct comparison with the combination treatment (PAS, purple), isolated treatment controls were included in every assay: column 11 contained bacteria with antibiotic alone (teal) whereas row H contained bacteria with phage alone (light purple). In addition to the growth (no treatment, green) controls, column 12 contained all sterility/blank controls (no bacteria: growth media alone - yellow, with antibiotic – pale blue, or with phage – pink). The red box highlights where phage is paired with sub-minimum inhibitory concentrations of antibiotic. The upper blue box represents the antibiotic-dominated killing region, and the lower purple box represents the phage-dominated killing region. Insets show representative growth curves of a synergistic interaction capturing OD_600_ values over 24 h for phage-antibiotic combination treatment (red) relative to the bacterial control (green), antibiotic treatment alone (blue), or phage treatment alone (purple) at the same concentrations and multiplicity of infection.

### Heatmaps Facilitate Efficient Visualization and Analysis of Phage-Antibiotic Interacting Regions

To quantify total growth over the 24-hour PAS assays (**Fig. 1**), area under the curve (AUC) was calculated for each well. Heatmaps were generated as colorimetric visualizations of these values with each well normalized to the untreated controls to facilitate cross-experimental comparisons. In the absence of perturbation, *P. aeruginosa* strain MPAO1pp-grew robustly reaching an AUC of 1508.9 ± 33.9 (average ± standard deviation) across assays. To identify the phage-antibiotic interacting regions, we first evaluated the effects of phage and antibiotic alone. In isolation, phage OMKO1 exhibited a 51.8 to 61.2% reduction across a multiplicity of infection ranging from 10^-7^ to 10 PFU/CFU (**Fig. 2A**). At an MOI of 10^-8^, phage efficacy was inconsistent across replicates. The initial titer of OMKO1 stock was ∼ 10^10^ PFU/mL. After several serial dilutions, the concentration of phage in column 11 was 4 PFU/mL, introducing ∼1.2 phage particles to a well containing 300 µL total. At such a low concentration, there may not have been phages introduced into the system, or there were not enough phages to encounter, infect, and replicate within host cells. Therefore, data collected at an MOI of 10^-8^ are shown in heatmaps for visualization purposes but excluded from synergy score calculations. The addition of antibiotics in isolation resulted in varying levels of reduction. Ciprofloxacin proved most efficient in inhibiting MPAO1pp-growth. At 0.5 µg/mL, ciprofloxacin achieved 93.17 ± 0.5% reduction. Aztreonam showed the least efficacy, requiring higher concentration to inhibit growth: at 8 µg/mL, aztreonam reduced growth by 83.38 ± 4.8%. Individual antibiotics achieved 80% growth inhibition (MIC_80_, determined by relative AUC) at the following concentrations: ciprofloxacin = 0.125 µg/mL (**Fig. 2B**), colistin = 2 µg/mL (**Fig. 2C**), tobramycin = 2 µg/mL (**Fig. 2D**), ceftazidime > 4 µg/mL (**Fig. 2E**), and aztreonam = 8 µg/mL) (**Fig. 2F**).

**Figure 2.**
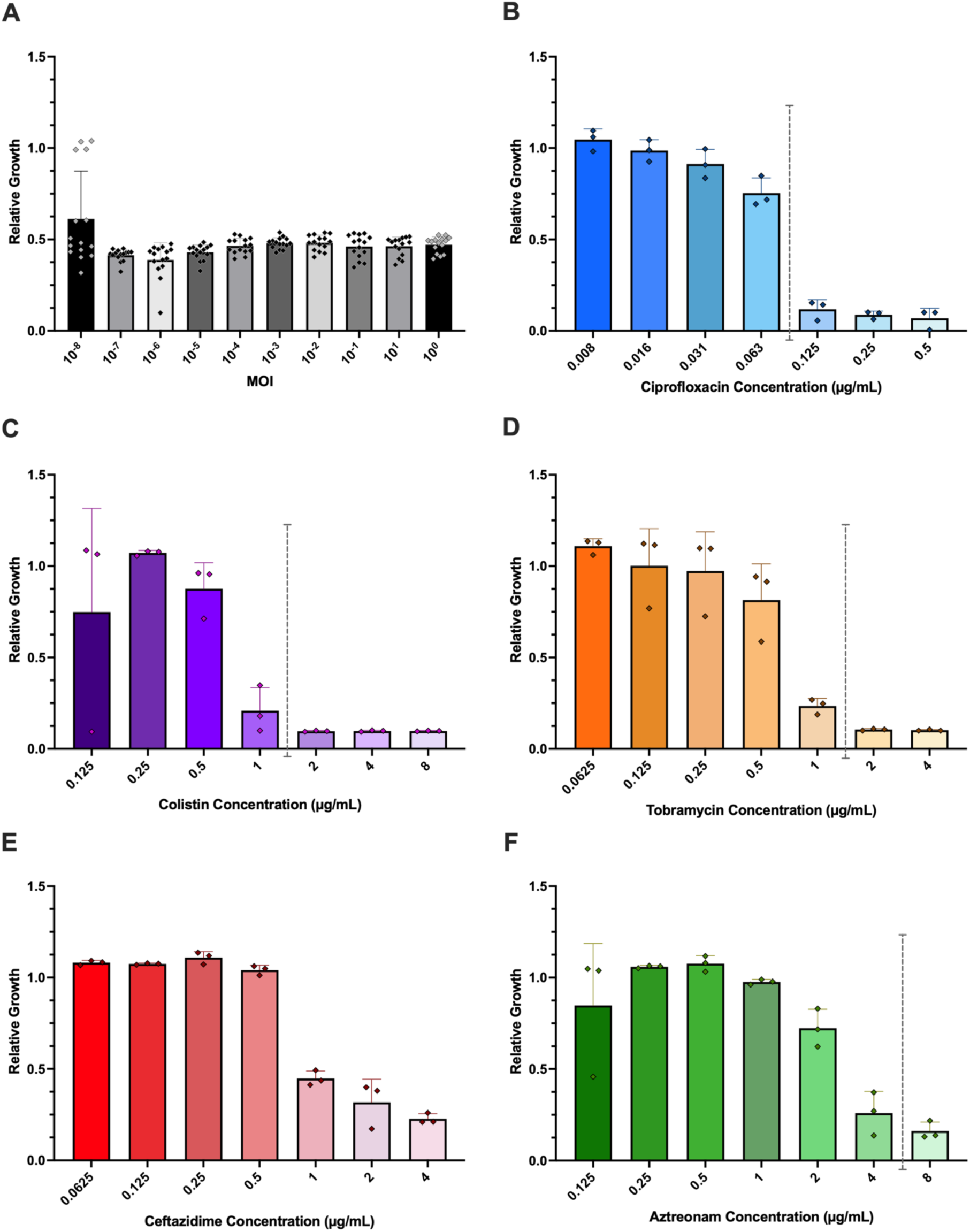
Phage and antibiotics alone inhibit bacterial growth to varying extents in isolation. The relative growth (AUC _phage alone or antibiotic alone_ / AUC _untreated growth control_) of strain MPAO1pp-in the presence of phage or antibiotics in isolation across (A) ten phage MOIs ranging from 10^0^ to 10^-8^ (n=15) or (B – F) five antibiotics at varying antibiotic concentrations (n=3 per antibiotic). Antibiotics are ordered B through F by descending efficacy. MIC_80_ displayed as vertical dashed line. AUC calculated from 24 h growth curve assays with OD_600_ readings taken every 10 minutes. Error bars represent standard deviation.

The antibiotic-dominated, phage-antibiotic interacting, and phage-dominated regions varied for each plate depending on antibiotic inhibition. For our study, the antibiotic-dominated region is where the antibiotic alone achieves ≥ 80% inhibition. The phage-antibiotic interacting region starts when the antibiotic alone inhibits < 80% bacterial growth and extends to the phage-dominated region, where the antibiotic reaches low concentrations that cease to reduce any growth. Regions were defined horizontally, not vertically, because bacterial reduction was comparable across phage MOIs (ranging from 51.84 – 61.24% reduction at concentrations within 10^0^ – 10^-7^ PFU/CFUs) (**Fig. 2A**). Therefore, the phage-antibiotic interacting region, or areas of potential synergy, additivity, or antagonism contained phage and antibiotics at wide ranges of sub-minimum inhibitory concentrations that were antibiotic-specific. Ciprofloxacin: 0.016 – 0.063 µg/mL (**Fig. 3A**); aztreonam: 0.5 – 4 µg/mL (**Fig. 3B**); ceftazidime: 0.5 – 4 µg/mL (**Fig. 3C**); tobramycin: 0.25 – 1 µg/mL (**Fig. 3D**); and colistin: 0.25 – 1 µg/mL (**Fig. 3E**). Heatmaps provide visualizations of the differences in bacterial growth observed between treatments: antibiotic alone, phage-antibiotic combination, and phage-alone. Phage-antibiotic combination wells presented as lighter colors on the heatmap when compared to respective antibiotic-only and phage-only controls, indicating less bacterial growth in combination. Ciprofloxacin alone at 0.063 µg/mL reduced total growth by 24.7% ± 8.3 (**Fig. 2B**). In comparison, ciprofloxacin at the same concentration reduced total growth by 91.7% ± 5.3 when paired with OMKO1 at 10^-2^ PFU/CFU (**Fig. 3A**). In isolation, 2 µg/mL aztreonam reduced growth by 27.6 ± 10.4% (**Fig. 2C**). When paired with OMKO1 at 10^-1^ PFU/CFU aztreonam (2 µg/mL) reduced growth by 89.7 ± 0.5% (**Fig. 3B**). Ceftazidime alone at 1 µg/mL reduced growth by 55.2 ± 4.1% (**Fig. 2E**). At the same concentration, ceftazidime paired with OMKO1 (10^-6^ PFU/CFU) reduced growth by 86.4 ± 0.4% (**Fig. 3C**). At 1.0 µg/mL, tobramycin reduced growth by 76.57 ± 4.19% (**Fig. 2D**). When paired with OMKO1 (10^-1^ PFU/mL), the combination reduced growth by 89.34 ± 0.39% (**Fig. 3D**). At 0.5 µg/mL, colistin inhibited growth by 12.38 ± 14.22% (**Fig. 2C**). When paired with OMKO1 (10^-5^ PFU/CFU), bacterial growth was reduced by 90.01 ± 0.36% (**Fig. 3E**). Collectively, we observed greater reductions in growth within the phage-antibiotic interaction region than either phage or antibiotic in isolation.

**Figure 3.**
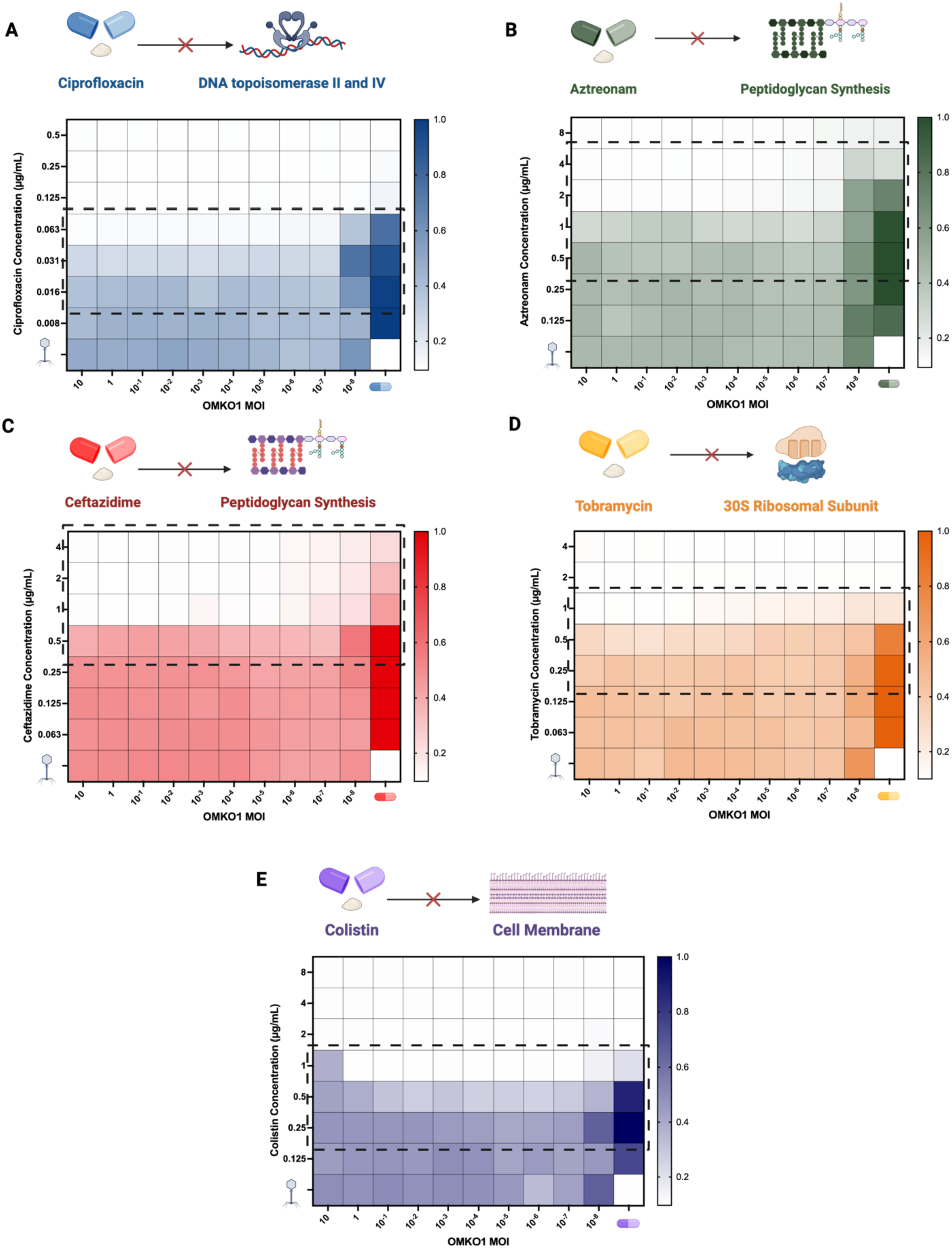
Heatmaps provide colorimetric visualization of phage-antibiotic synergy assays that aide in identification of regions with phage-antibiotic interactions. The effects of phage-antibiotic combination treatments for phage OMKO1 with five antibiotics from diverse drug classes are shown A) Ciprofloxacin, fluoroquinolone; B) Aztreonam, monobactam; C) Tobramycin, aminoglycoside; D) Ceftazidime, cephalosporin; E) Colistin, polymyxin. Heatmaps represent the mean “total growth” of three biological replicates over 24 h calculated as follows: Total growth = 1 – [(AUC _treated well_) / (AUC _untreated control well_)]. Each heatmap is normalized to column 12 untreated growth controls, shown as 1.0 or 100% growth in the corresponding color bars. White represents high bacterial killing (>90% reduction in growth). Phage-antibiotic interacting regions are boxed by dashed lines. Graphics above heatmaps created in https://BioRender.com.

### The Bliss Independence Model of Synergy Classifies Phage-Antibiotic Interactions

To quantify synergy, additivity, or antagonism for each treatment combination, the Bliss independence model was applied to the AUC data from the PAS assays, normalized to the untreated growth controls. Specifically, the observed effect on growth for a given treatment was calculated as a proportion of the untreated controls (i.e. R_treatment_ = AUC_treatment_/AUC_control_) and used as input into the model. The Bliss independence model determines synergy (ΔR) by comparing the observed effect of a combination treatment (R_abx, Φ_) with the expected effect of a combination treatment (33). The expected effect is the sum of the two treatment effects observed in isolation (R_abx_ + R_Φ_). This model was used for these combinations due to the distinct mechanisms of action (MOA) across the panel of antibiotics and the assumption that these MOAs differ from that of lytic phage OMKO1. The null hypothesis is that the reduction_observed_ – reduction_expected_ is equal to zero, indicating there is no advantage to using the treatments together rather than independently. If the reduction_observed_ – reduction_expected_ is less than zero, the combination is deemed antagonistic. Finally, if the reduction_observed_ – reduction_expected_ is greater than zero, the combination is termed synergistic: using the treatments together is more effective than the sum of either agent used independently (**Fig. 4A**). By this model, ΔR values range (–1.0 to 1.0), with –1.0 being 100% antagonistic, 0.0 being additive, and +1.0 being 100% synergistic. If a ΔR value is 0.25, this indicates that the tested combination inhibited 25% more bacterial growth than expected. Importantly, the Bliss independence model accounts for the fact that a cell can only be killed once (33). It does this by subtracting the product of the reduction by antibiotic (R_abx_) and reduction by phage (R_Φ_) from their sum (**Fig. 4A**). This is a notable feature of the Bliss independence model which distinguishes it from other commonly used models for determining synergy (34).

**Figure 4.**
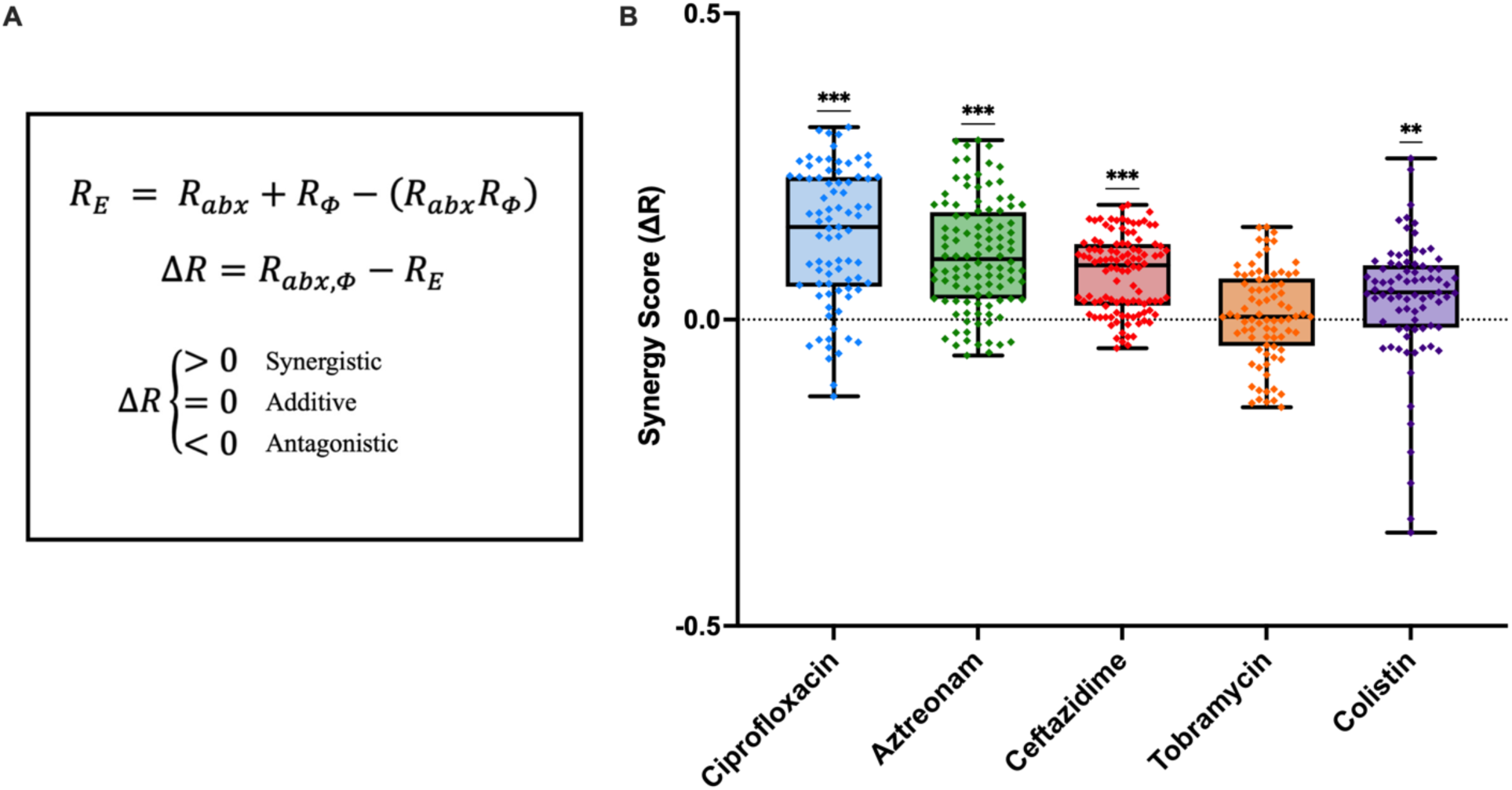
Phage OMKO1 synergizes with ciprofloxacin, aztreonam, ceftazidime, and colistin based on synergy score determined by the Bliss Independence Model. (A) Bliss Synergy Score calculation. R_E_: expected reduction by combination treatment, R_abx_: bacterial inhibition by antibiotic alone (AUC_abx_/AUC_control_), R_Φ_: bacterial inhibition by phage alone (AUC_Φ_/AUC_control_), R_abx, Φ_: actual (observed) reduction by combination treatment (AUC_abx,Φ_/AUC_control_). R_E_ is calculated by subtracting the product of R_abx_ and R_Φ_ from their sum. ΔR is subsequently calculated by subtracting this value from R_abx, Φ_. The combination is considered synergistic when ΔR > 0, additive when ΔR = 0, and antagonistic when ΔR < 0. (B) Synergy scores within each assay’s phage-antibiotic interactive zone. Plot represents mean synergy scores across varying antibiotic concentrations and phage multiplicities of infection. Ciprofloxacin = 0.016 – 0.063 µg/mL; aztreonam = 0.5 – 4 µg/mL; ceftazidime = 0.5 – 4 µg/mL; tobramycin = 0.25 – 1 µg/mL; and colistin = 0.25 – 1 µg/mL, each including intermediate 0.5 intervals. MOI ranges from 10^-7^ – 10 PFU/CFU. Each concentration combination was performed in biological triplicate. Each dot represents one replicate of a phage-antibiotic combination. Line within boxes positioned at the mean, with whiskers spanning the minimum to maximum values. Individual data points are shown in Supplemental Table 1. A one sample t test and Wilcoxon test with a hypothetical value of zero were performed to measure statistical significance. Significance thresholds: P > 0.12, not significant (ns); P* < 0.033; P** < 0.002; P*** < 0.001. Same data as figure 3.

To consider synergy robust to variations in phage and antibiotic concentrations that may be achieved in vivo, we first considered overall synergy across all phage MOIs and antibiotic concentrations within phage-antibiotic interactive regions. Ciprofloxacin (fluoroquinolone, target: DNA topoisomerase II & IV), aztreonam (monobactam, target: Penicillin-Binding Protein-3, PBP-3), ceftazidime (cephalosporin, target: PBP-3), and colistin (polymyxin, target: outer-membrane lipopolysaccharide (LPS) molecules) achieved average synergy scores > zero across phage MOIs (10^-7^ PFU/CFU –10 PFU/CFU) and antibiotic concentrations within their respective ranges. The synergy scores for tobramycin (aminoglycoside, target: 30S Ribosomal Subunit) paired with OMKO1 ranged from -0.078 to 0.151 and did not show a statistically significant difference from zero, indicating the average combined effect of tobramycin with phage OMKO1 is additive (**Fig. 4B**).

Ciprofloxacin-OMKO1 combinations achieved an average synergy score of 0.135 ± 0.111 (ΔR ± standard deviation) with a lower – upper 95% confidence interval (CI_95_) of 0.111 – 0.160 (**Fig. 4B**). The maximum ΔR value of 0.314 ± 0.045 occurred at a ciprofloxacin concentration of 0.063 µg/mL and phage MOI of 10^-3^ PFU/mL. The expected bacterial growth reduction of this combination was 64.99%. The observed reduction of this combination was 96.41%. Aztreonam-OMKO1 combinations achieved a comparable mean synergy score of 0.105 ± 0.088 (CI_95_ = 0.088 – 0.121), with a maximum ΔR of 0.293 ± 0.057 at an aztreonam concentration of 2.0 µg/mL and an OMKO1 MOI of 10^-3^ PFU/mL (**Fig. 4B**). The expected reduction of this combination was 60.87%; the observed reduction was 90.17%. Ceftazidime-OMKO1 combination synergy scores averaged at 0.075 ± 0.622 (CI_95_ = 0.063 – 0.087) (**Fig. 4B**). The highest ΔR, 0.187, occurred at a ceftazidime concentration of 0.5 µg/mL and an OMKO1 MOI of 10 PFU/mL. This combination achieved 18.72% more reduction than expected. The average synergy score for colistin-OMKO1 combinations was 0.032 ± 0.104 (CI_95_ = 0.008 – 0.054). ΔR scores for OMKO1-colistin combinations spanned a greater range than the other combinations but achieved a maximum synergy score of 0.263 (colistin = 0.5 µg/mL; OMKO1 = 10^-2^) (**Fig. 4B**). On average, tobramycin-OMKO1 combinations were additive with a mean synergy score of 0.006 ± 0.075 (CI_95_ = -0.011 – 0.023) (**Fig. 4B**).

### Antibiotic concentration impacts synergy scores

While synergistic effects were observed in data compiled across antibiotic concentrations and different multiplicities of infection for phage OMKO1 (**Fig. 4B**), there was a fair amount of spread apparent in the data across all tested antibiotics. To begin to parse these effects, we sought to assess the Bliss metric of synergy between antibiotic concentrations individually. Synergy within phage-antibiotic interactive zones varied depending on the antibiotic concentration (**Fig. 5**). OMKO1-ciprofloxacin combinations were synergistic across all three antibiotic concentrations, achieving a maximum synergy score of 0.314 at 0.063 µg/mL; this combination reduced growth by 96.41%, notably higher than its expected reduction of 64.99% (**Fig. 5A**). OMKO1-aztreonam combinations were synergistic at antibiotic concentrations between 0.5 and 2 µg/mL, with a maximum synergy score of 0.293 at 2 µg/mL. At 4 µg/mL, OMKO1-aztreonam combination effects did not show a statistically significant difference from zero, indicating additivity (**Fig. 5B**). OMKO1-ceftazidime combinations were synergistic between 0.5 and 2 µg/mL. The pairing was most synergistic at 0.5 and 1 µg/mL and least synergistic at 2 µg/mL (**Fig. 5C**). At 4µg/mL, phage-ceftazidime combinations were additive; they did not show a statistically significant difference from zero (**Fig. 5C**). Phage-tobramycin combinations showed an overall decrease in efficacy compared to other phage-antibiotic combinations (**Fig. 5D**). Phage-tobramycin combinations at 1 µg/mL showed a statistically significant mean negative ΔR, indicating antagonism at this concentration. However, as tobramycin concentration decreased, ΔR values increased: at 0.25 µg/mL and 0.5 µg/mL tobramycin with OMKO1 achieved mean synergy scores above zero (**Fig. 5D**). OMKO1-colistin combinations were synergistic at concentrations from 0.25 to 0.5 µg/mL with a maximum synergy score occurring at a colistin concentration of 0.5 µg/mL (ΔR = 0.263) (**Fig. 5E**). This combination reduced growth by 89.84%, which was 26.31% more effective than the expected reduction (63.53%). At 1 µg/mL, colistin-OMKO1 combinations were, on average, additive.

**Figure 5.**
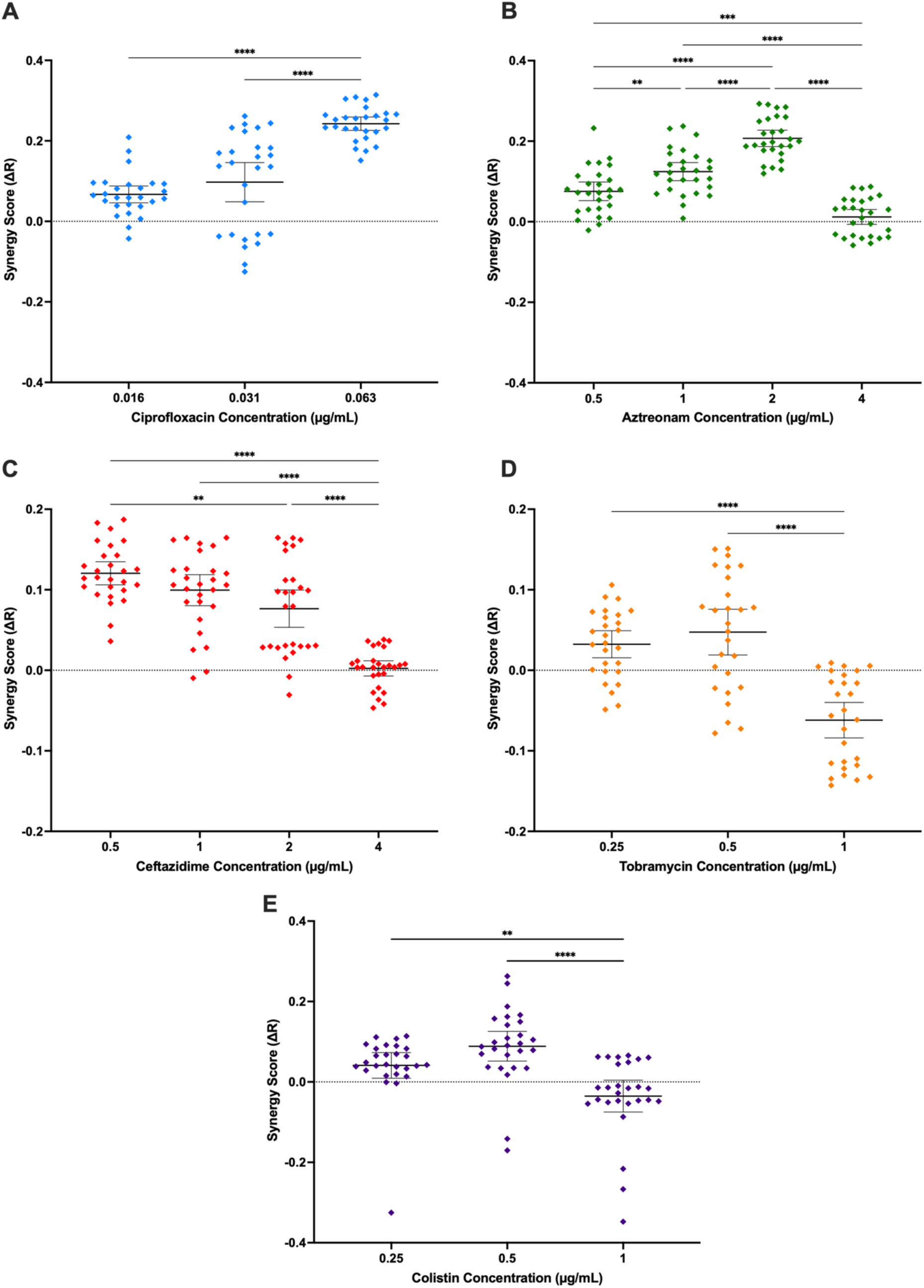
Synergy scores within each phage-antibiotic interacting zone vary by antibiotic concentration. ΔR values of the Bliss Independence Model were calculated using AUC data from 24 h growth curve assays. Antibiotic concentration ranges varied: (A) ciprofloxacin from 0.016 to 0.063 µg/mL; (B) aztreonam from 0.5 to 4 µg/mL; (C) ceftazidime from 0.5 to 4 µg/mL; (D) tobramycin from 0.25 to 1 µg/mL; and (E) colistin from 0.25 to 1 µg/mL. All assays performed in biological triplicate. Lines represent the mean value with error bars displaying standard deviation. Ordinary one-way analysis of variance (ANOVA) and Tukey’s multiple comparisons test were used to determine statistically significant differences between antibiotic concentrations. Significance thresholds P > 0.12, not significant (ns); P* < 0.033; P** < 0.002; P*** < 0.001. Same input data as Figure 4.

**Figure 6.**
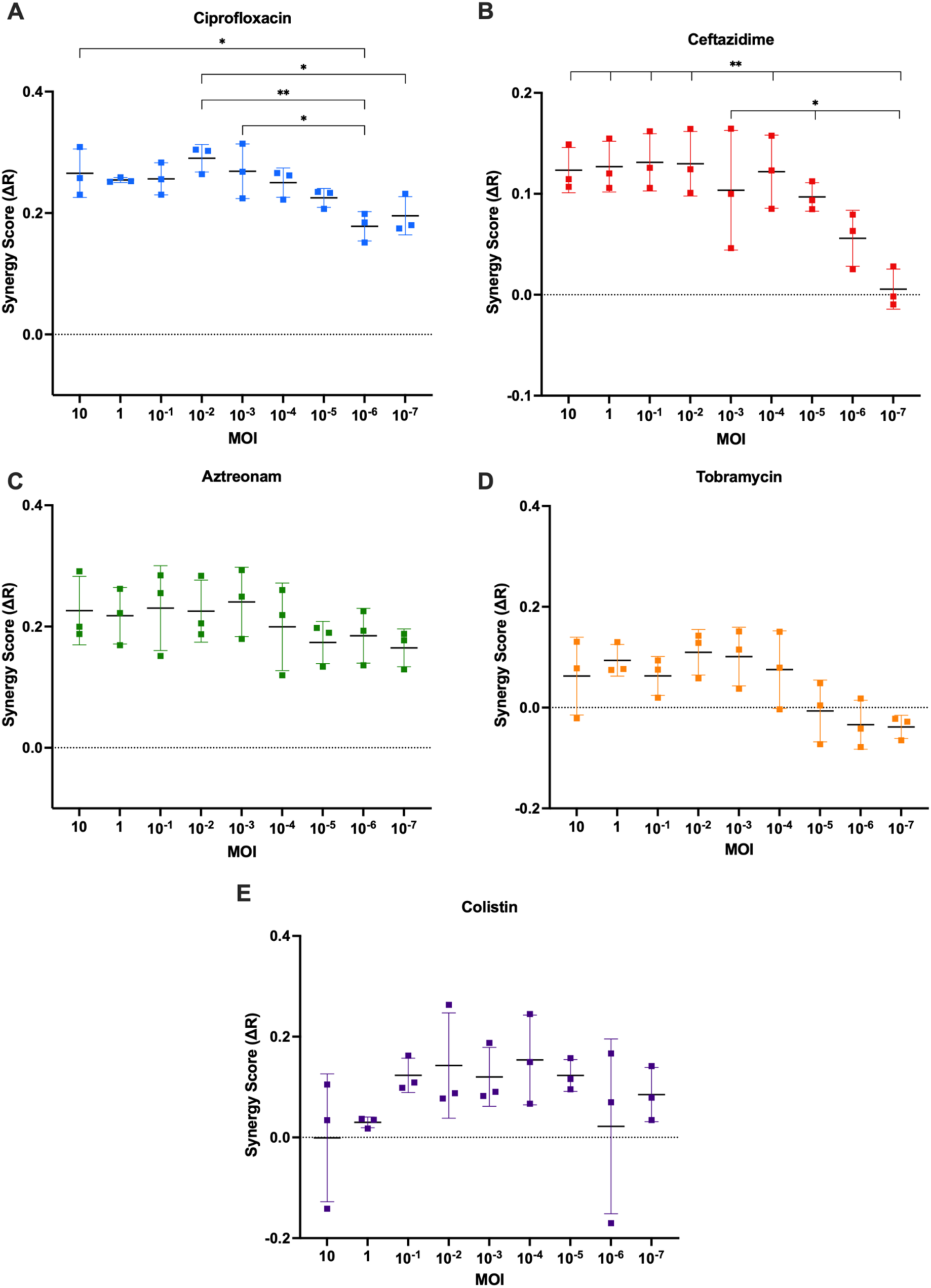
Effects of phage MOI on Bliss synergy scores vary by antibiotic. Bliss synergy scores were calculated for five antibiotics at sub-minimum inhibitory concentrations and a range of phage MOIs (10^-7^ PFU/CFU – 10 PFU/CFU). Antibiotics were assessed at the following concentrations: Ciprofloxacin = 0.063 µg/mL; aztreonam = 2.0 µg/mL; ceftazidime = 1.0 µg/mL; tobramycin = 0.5µg/mL; colistin = 0.5 µg/mL. Each graph represents the mean of three biological replicates with error bars capturing standard deviation. Statistical significance was measured with an ordinary one-way analysis of variance (ANOVA) and Tukey’s multiple comparisons test. Significance thresholds: P > 0.12, not significant (ns); P* < 0.033; P** < 0.002. Same data shown in figure 4

### Comparing Bliss synergy score of varying phage MOIs and constant antibiotic concentration

After analyzing how antibiotic concentration impacts phage-antibiotic combination treatment success, synergy scores were calculated for each antibiotic at a constant, sub-minimum inhibitory concentration while phage was varied across nine MOIs (**Fig. 7**). Ciprofloxacin-OMKO1 combinations showed lower synergy scores at lower MOIs compared to higher ones. The lowest synergy score, ΔR = 0.151, occurred at an MOI of 10^-6^ PFU/CFU. This score was lower than those achieved at higher MOIs: 10 PFU/CFU (0.266 ± 0.039; ΔR ± standard deviation), 10^-2^ PFU/CFU (0.290 ± 0.023), and 10^-3^ PFU/CFU (0.269 ± 0.045). Similarly, at an MOI of 10^-7^, the combination was less synergistic (ΔR = 0.195) than it was at an MOI of 10^-2^ (ΔR = 0.29). Despite the variation across MOIs, values remained positive (**Fig. 7A**). Ceftazidime also showed differences in synergy scores at a low MOI compared to higher ones. At 10^-7^ PFU/CFU, ΔR = 0.006 ± 0.0199. This score was notably lower than those achieved at MOIs between 10 and 10^-5^ PFU/CFU, which ranged 0.046 – 0.165 (**Fig. 7C**). Aztreonam, tobramycin, and colistin showed no significant difference in synergy at any of the nine MOIs ranging from 10 PFU/CFU to 10^-7^ PFU/CFU (**Fig. 7C, D, & E**). At 0.5 µg/mL, synergy scores of aztreonam-OMKO1 combinations spanned 0.174 – 0.241. Scores remained positive and comparable across MOIs (**Fig. 7C**). Tobramycin-OMKO1 combinations were additive at 0.5 µg/mL and did not vary across MOIs (**Fig. 7D**).

**Figure 7.**
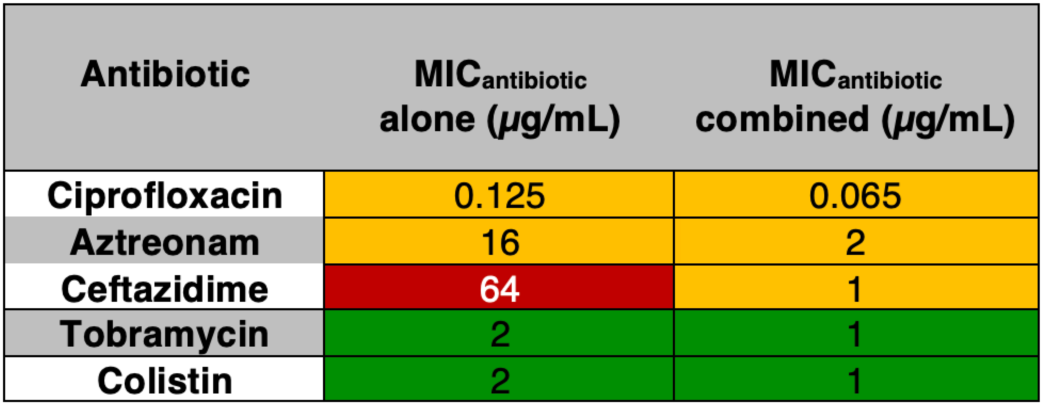
Phage-antibiotic combinations reduce the relative MIC of five distinct antibiotic classes. MICs determined with liquid growth assays by exposing bacteria to varying concentrations of antibiotic (n=3). MIC defined as the smallest concentration required to achieve no visible turbidity after 24 hours of growth in liquid. Entries colored according to EUCAST clinical breakpoint data: susceptible (green), intermediate (yellow), and resistant (red).

### All tested phage-antibiotic combination treatments reduce MIC by ≥ 2-fold

Regardless of Bliss synergy score, when combined with phage OMKO1, all five tested antibiotics demonstrated a reduction in minimum inhibitory concentration (MIC). MIC_80_ values, used to define regions within the PAS assays, were determined using relative growth values. Because not all PAS assays extended beyond MIC_80_, additional assays were performed separately with more expansive ranges of antibiotic concentrations to obtain MICs, defined as the lowest concentration of antibiotic required to achieve no visible turbidity of a liquid culture following 24 hours of incubation. These values were then compared to antibiotic concentrations of combination treatment wells within the phage-antibiotic interacting zones that were clear after 24 hours. Phage OMKO1 (10^-1^ PFU/CFU) with ciprofloxacin reduced the MIC by two, aztreonam by eight, ceftazidime by 64, tobramycin by two, and colistin by two (**Fig. 7**). Based on clinical breakpoint data (35), the reduction of ceftazidime MIC from 64 µg/mL to 1 µg/mL re-classified strain MPAO1pp-from resistant to intermediate.

### Combination treatments limit resistant regrowth over 24 hours

In addition to an overall reduction in antibiotic MIC, phage-antibiotic combination treatments extended the suppression of resistant regrowth across all tested antibiotics, regardless of synergy score and antibiotic class. Despite an initial reduction in growth across all five antibiotic treatments, the inhibition was eventually overtaken by growth of resistant bacteria which was particularly evident when applied alone at sub-minimum inhibitory concentrations with growth rebounding between 2.8 and 19.5 hours depending on the antibiotic (**Fig. 8A**). A similar growth pattern indicative of resistance was observed at all phage MOIs ranging from 10 PFU/CFU to 10^-7^ PFU/CFU. On average, this regrowth occurred at 11 hours (**Fig. 8B**). Additional analyses of wells within phage-antibiotic interacting zones were performed to determine how phage-antibiotic combinations may affect regrowth of resistant bacteria over 24-hours. All five phage-antibiotic combination treatments show a reduction in resistant regrowth compared to their respective individual treatments as seen by their 24-hour growth curves (**Fig. 8C–G**). Resistant regrowth time is most dramatically affected by the addition of OMKO1 to ciprofloxacin and aztreonam (**Fig. 8H**). At 0.063 µg/mL, ciprofloxacin suppressed bacterial growth for 4.867 ± 0.115 hours. When combined with OMKO1 (10^-2^ PFU/CFU), resistant regrowth was suppressed for the full 24-hour incubation (**Fig. 8C & H**). Aztreonam inhibited growth for 3.367 ± 0.115 hours alone at 2 µg/mL. With the addition of OMKO1 (10^-3^ PFU/CFU), bacterial suppression lasted for 22.733 ± 2.194 hours (**Fig. 8D & H**). Ceftazidime-alone, phage-alone, and phage-ceftazidime combination treatments share similar growth curves for the first 3.8 hours. At this time point, bacteria treated with only ceftazidime continue to grow until around 8 hours, when they reach a plateau. At 3.8 hours, the phage-only curve shows OMKO1 (10^-6^ PFU/CFU) infecting and killing bacteria, resulting in a drop in growth until resistance arises at 11.133 ± 0.764 hours. Ceftazidime-only curves show a second rise in growth after 20.133 ± 0.289 hours. In contrast, the combined treatment suppresses growth for the entire 24 hours (**Fig. 8E & H**). Tobramycin shares similar differences in the suppression time achieved by combination treatment vs. tobramycin-alone treatment. At 1 µg/mL, tobramycin suppresses bacterial growth for 17.3 ± 1.323 hours. When paired with OMKO1 (10^-2^ PFU/CFU), resistant regrowth is suppressed for 24 hours (**Fig. 8F & H**). At 1 µg/mL, colistin suppresses bacterial growth for 18.233 ± 4.050 hours. With the addition of OMKO1 (10^-5^ PFU/CFU), resistant regrowth is suppressed for the entire 24-hour period (**Fig. 8G & H**).

**Figure 8.**
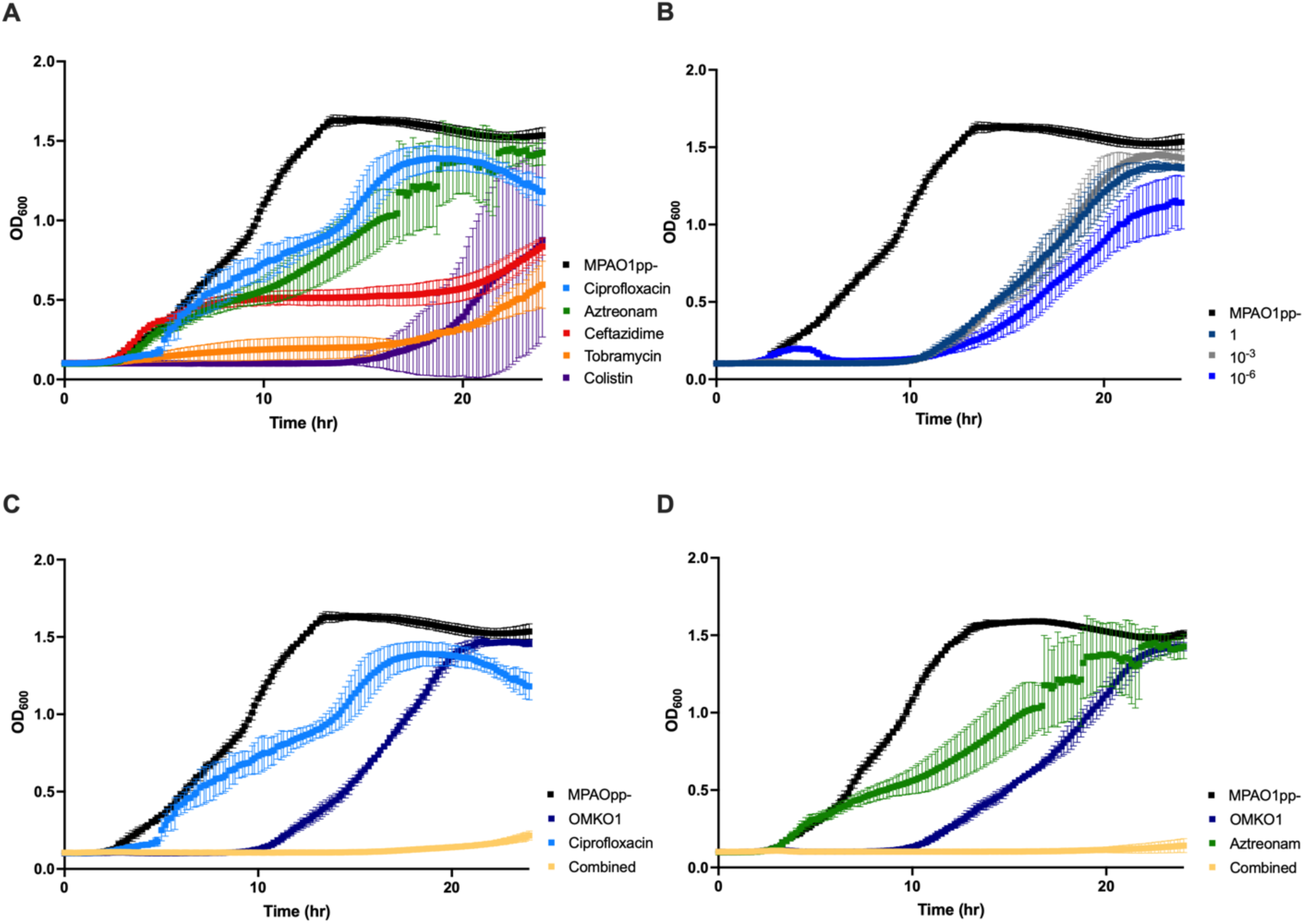
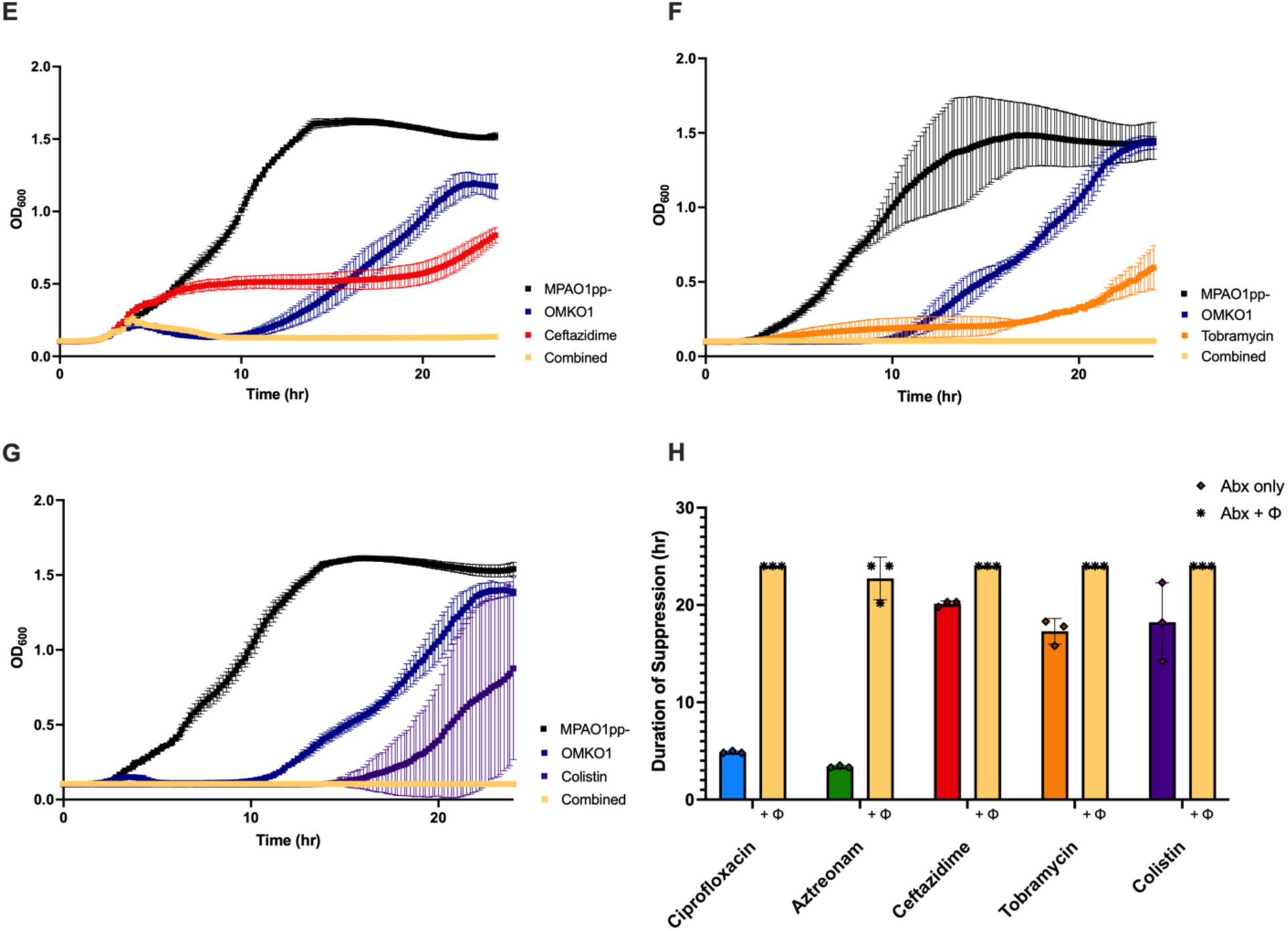
Growth curves of MPAO1pp-at varying antibiotic concentrations, phage MOIs, and phage-antibiotic combination treatments. Bar graph representing time that resistant regrowth is suppressed by antibiotic-alone vs. combination treatments. (A) Growth curves of MPAO1pp-at sub-minimum inhibitory concentrations of five antibiotics: ciprofloxacin = 0.063 µg/mL, aztreonam = 2 µg/mL, ceftazidime = 1 µg/mL, tobramycin = 1 µg/mL, and colistin = 1 µg/mL. (B) Growth curves of MPAO1pp-when exposed to OMKO1 at MOIs of 1 PFU/CFU, 10^-3^ PFU/CFU, and 10^-6^ PFU/CFU. (C–G) Growth curves of MPAO1pp-at sub-minimum inhibitory concentrations of antibiotic and phage. Antibiotic and phage concentrations vary per combination: ciprofloxacin = 0.063 µg/mL, MOI = 10^-2^ PFU/CFU; aztreonam = 2 µg/mL, MOI = 10^-3^ PFU/CFU; ceftazidime = 1 µg/mL, MOI = 10^-6^ PFU/CFU; tobramycin = 1 µg/mL, MOI = 10^-2^ PFU/CFU; and colistin = 1 µg/mL, MOI = 10^-5^ PFU/CFU. Error bars represent standard deviation. Area under curve data and growth curves generated from OD_600_ readings taken every 10 minutes (n=3). (H) Bar graph comparing amount of time combination treatments suppress resistant regrowth to amount of time regrowth is suppressed by corresponding antibiotic-alone treatments. Time of regrowth determined by inflection points on growth curves. Error bars represent standard deviation (n=3). Same data shown in figure 3.

## Discussion

### Effects of antibiotic class on Phage-Antibiotic Synergy (PAS)

Previous studies have suggested that antibiotic class and mechanism of action may predict phage-antibiotic interactions, and therefore, PAS or antagonism (25). These studies have been driven by the assumption that antibiotics like macrolides, fluoroquinolones, or aminoglycosides, may confer antagonism when paired with bacteriophages, since phages require host machinery for their survival and replication (26,29). Limiting this pertinent machinery could reasonably impede the phage’s ability to replicate within the host and lyse. In our study, these antibiotics are: ciprofloxacin, a fluoroquinolone, and tobramycin, an aminoglycoside (36,37). PAS assays with these antibiotics revealed that tobramycin-OMKO1 pairings were additive. However, ciprofloxacin-OMKO1 combinations were highly synergistic. Despite these antibiotics’ similarities in mechanism of action, the differences in their specific targets could explain why ciprofloxacin conferred synergy while tobramycin did not. Ciprofloxacin inhibits bacterial type II topoisomerases (DNA gyrase and topoisomerase IV), enzymes primarily responsible for limiting DNA supercoiling during replication (38). In comparison, tobramycin inhibits bacterial growth by targeting the 30S ribosomal subunit. It is possible that phage OMKO1, a T-even-like phage, encodes its own bacteriophage topoisomerase II, thus compensating for compromised bacterial DNA gyrase in the presence of ciprofloxacin (39).

Independently encoding DNA topoisomerase, as phage T4 and T6 do, could also explain ciprofloxacin-OMKO1 combinations’ continued rise in synergy score even at higher antibiotic concentrations. In contrast, there are not currently any phage known to encode their own ribosomal machinery, resulting in a more significant hindrance of OMKO1 efficiency in the presence of ribosomal subunit-targeting antibiotics like tobramycin. These findings suggest that bacteriophage productivity may be more significantly impacted by translation-targeting antibiotics than they are by transcription-targeting ones, emphasizing the significance of cross-checking antibiotic-targets with specific phage genomes when designing combination therapies.

The other three antibiotics tested in our study: aztreonam (a monobactam), ceftazidime (a cephalosporin), and colistin (a polymyxin), are cell envelope-active agents (40–42). Previous studies have suggested that these antibiotics would result in synergy when paired with phage, because antibiotic classes which disrupt bacterial cell wall synthesis or stability may synergistically enhance phage’s ability to lyse the bacterial cell following replication (25). While our findings show synergy across these antibiotics, the previously noted synergy seen by ciprofloxacin-phage combinations suggest that antibiotic class alone does not consistently predict phage-antibiotic synergy or antagonism (**Fig. 4B**), and that PAS may be better predicted by comprehensive considerations of conditions including bacterial strain, antibiotic target, and phage type. These conclusions align with those in Coyne et al.’s study comparing protein synthesis-targeting antibiotics with cell envelope active ones (29).

While PAS may not be predictable based solely on antibiotic class, our findings display a correlation between phage-antibiotic synergy and antibiotic-induced filament formation. In our study, the three combinations yielding highest average synergy scores were ciprofloxacin-OMKO1, aztreonam-OMKO1, and ceftazidime-OMKO1, three filamentation-inducing antibiotics (43–45). In a recent study, Bulssico et al. investigate the effects of sub-minimum inhibitory concentrations of filamentation-inducing antibiotics (including ceftazidime and ciprofloxacin) on gram-negative bacteria and how these effects influence PAS (46). Sub-lethal doses of these antibiotics prompted stress tolerance and DNA repair pathways, including the inhibition of bacterial division and antibiotic-induced filamentation. This filamentation resulted in enlarged bacterial surface area, and in turn, increased phage adsorption and burst size; specifically, adsorption per cell increased two-fold in bacteria treated with sub-lethal concentrations of ciprofloxacin compared to untreated bacteria (46). Given these findings, we hypothesize that the high synergy observed between phage and ciprofloxacin, aztreonam, and ceftazidime may be a result of these antibiotics inducing filamentation of the bacteria, which enlarges cell surface area, and increases the probability of phage adsorption, infection, and lysis. These results highlight the significance of considering antibiotic-induced morphology changes in phage-combination therapy design and of studying these interactions on both a population scale and individual one.

### Effects of antibiotic concentration and MOI on phage-antibiotic synergy and antagonism

In addition to antibiotic class, we evaluated how antibiotic concentration may impact phage-antibiotic synergy by comparing scores within phage-antibiotic interacting zones (**Fig. 5**). All antibiotic-phage combinations displayed their lowest synergy scores at the highest, sub-minimum inhibitory concentration of antibiotic, except for ciprofloxacin. Ciprofloxacin-OMKO1 synergy scores uniquely increased with antibiotic concentration. We propose that this trend may be explained by ciprofloxacin specific-induced changes in bacterial gene expression. One study that investigated proteomic alterations of ciprofloxacin-exposed *P. aeruginosa* noted an increase in the expression of efflux pump MexAB-OprM, a putative target of phage OMKO1 (45,47). By upregulating efflux pumps, ciprofloxacin exposure increases the probability of phage encountering a target and successfully adsorbing to and replicating within bacteria. This explanation also highlights the potential of phage-steering, or intentionally pairing antibiotics and phage with complementary effects: as bacteria experience an upregulation in efflux pumps, becoming resistant to ciprofloxacin, they are simultaneously overexpressing a proposed target of OMKO1, becoming more susceptible to phage (47). This idea is speculative, as the possible receptor-binding target(s) for phage OMKO1 remain the focus of continued investigation (48,49).

Aztreonam-OMKO1 synergy scores also increased with antibiotic concentration from 0.5 – 2 µg/mL before falling at 4 µg/mL (**Fig. 5B**). Ceftazidime-OMKO1 synergy scores showed a downward trend in synergy scores as antibiotic concentration increased, reaching a minimum average ΔR at its highest, sub-minimum inhibitory concentration (4 µg/mL) (**Fig. 5C**). Interestingly, previous literature has highlighted a dose-dependent relationship between antibiotic-induced filamentation and antibiotic concentration specifically with ceftazidime (43). In their study, Buijs et al. determine that at low ceftazidime concentrations, PBP-3 inhibition results in filament formation; however, at higher concentrations of ceftazidime, inhibition of PBP-1 rapidly lyses bacteria (43). If antibiotic-induced filamentation is, in fact, a key driver of PAS, this shift from ceftazidime-induced filamentation to ceftazidime-induced lysis could account for the dip in synergy scores at 4 µg/mL (**Fig. 5C**). This increase in cell lysis by ceftazidime at a higher concentration may lower OMKO1’s efficiency by limiting opportunities for the phage to adsorb to bacteria with intact cell walls, rendering their combination additive rather than synergistic. Aztreonam-OMKO1, ceftazidime-OMKO1, and colistin-OMKO1 combination treatments all shifted from synergistic to additive at their highest, sub-minimum inhibitory concentration of antibiotic (**Fig. 5 B, C, & E**). Tobramycin-OMKO1 shifted from synergistic to antagonistic at its highest, sub-minimum inhibitory concentration of antibiotic (**Fig. 5D**).

To assess the impact of phage concentration on PAS, synergy was evaluated at a constant, sub-minimum inhibitory concentration of antibiotics across MOIs from 10^-7^ PFU/CFU to 10 PFU/CFU. In our investigations, two of the antibiotic-phage pairings showed decreased synergy scores at lower MOIs: ciprofloxacin-OMKO1 and ceftazidime-OMKO1. Ciprofloxacin-OMKO1 combination treatments showed decreased synergy scores at MOIs below 10^-5^ PFU/CFU compared to 10^-4^ – 10 PFU/CFU (**Fig. 6A**). These findings align with those in a study by Namonyo et al. which applied phage to *P. aeruginosa* biofilm at varying MOIs and reported that at a low MOI, phage had reduced ability to inhibit bacterial growth (50). In our study, however, this was not a global trend across antibiotics. Outside of ciprofloxacin and ceftazidime, antibiotic-phage combinations were not significantly impacted by altering viral load. These findings indicate that PAS may lower the amount of phage required for bacterial treatments. Limiting the initial dose requirement is significant for cost and safety in the clinical setting as the phage will only replicate as they encounter susceptible bacteria (51).

### PAS reduces the minimum inhibitory concentration of all five tested antibiotics

PAS remained effective at sub-minimum inhibitory concentrations of all five antibiotics, effectively reducing their MIC (**Fig. 6**). Our findings support previous literature which also reports dramatic reductions of antibiotic MIC up to 32-fold when the antibiotic is paired with phages (32). MIC was most dramatically reduced for ceftazidime and aztreonam (**Fig. 7**), both of which are cell wall-active antibiotics that target PBP-3. This reduction in MIC is increasingly important when considering the rise of antimicrobial resistance, cost, and patient exposure to adverse side effects (52). First, reducing the initial dose of antibiotics during treatment can limit the overuse of antibiotics and therefore, antimicrobial resistance. Second, several antibiotics which have been traditionally viewed as “last-resort antibiotics,” including colistin, impose substantial side effects on patients (53). Unfortunately, these antibiotics are losing their antimicrobial efficacy due to increasing resistance, requiring higher dosages, prolonged treatment courses, and therefore, more severe side effects ranging from nephrotoxicity to neurotoxicity (54). PAS presents the potential to slow both the climbing levels of antimicrobial resistance and to improve the experience of patients who require long-term antibiotic usage by lowering necessary dosages.

### PAS limits the emergence of phage and antibiotic resistance

Since their initial discovery in 1917, interest in the development of phages for therapy has varied (55). In addition to the introduction of antibiotics, phages presented their own drawbacks which may have derailed their initial popularity, including their narrow host range and rapid development of evolved phage-resistance in target bacteria during treatment (56). It is true that when applied alone, phage OMKO1 encountered rapid emergence of resistance by *P. aeruginosa* (**Fig. 8B**). However, pairing OMKO1 with all five tested antibiotics resulted in decreased emergence of these resistant bacteria (**Fig. 8C–H**). This finding aligns with several other studies which highlight the ability of PAS to suppress the emergence of phage resistance (16,25,56–59). This capacity to limit phage resistance has been cited as a potential explanation for PAS in addition to: targeting different sites, re-sensitization of antibiotic-resistant bacteria, phage-mediated disruption of biofilms, and changes in bacterial cell morphology (60,61). Rationale for this phenomenon has been proposed from an evolutionary standpoint as well. Presenting two selective pressures on a system may limit the probability of the population undergoing beneficial mutations against both pressures. It is less likely for the bacteria to undergo random mutations that are beneficial against *both* the antibiotic and phage (56,62). This “tradeoff” hypothesis has been explored in previous studies that use phages to steer antibiotic re-sensitization (63,64).

Our study also presents limitations that should be addressed in future investigations. Firstly, PAS here was studied only using simultaneous treatment schedules. Several studies have demonstrated that simultaneously applying separate selective pressures to the bacteria is more effective in suppressing the development of phage resistance than applying them sequentially; however, compassionate usage of phage therapy largely occurs after a patient has been on antibiotics, not before (56,65,66). Additional studies should explore PAS when phage is added to antibiotic-resistant bacteria with that antibiotic present in the system. To render these studies more clinically relevant, similar experiments should also be performed with clinical strains. Our findings indicate that *P. aeruginosa* typically developed resistance to phage OMKO1 after ∼11 hours (**Fig. 8B**). Therefore, future studies should also investigate sequential treatment schedules with the addition of antibiotic at this time point with resistance profiling occurring pre- and post-treatment. Another limitation of this study is that differences in phage dynamics were not monitored across the five different antibiotics. Future studies should explore how antibiotic presence affects the lytic phage cycle, specifically the latent period and burst size. Lastly, this study focused on a set of clinically relevant, bactericidal antibiotics. Future studies should expand this set to include bacteriostatic antibiotics since bactericidal and bacteriostatic antibiotics’ differing impacts on bacterial metabolism may affect phage efficacy and PAS (29).

In conclusion, this study demonstrates the beneficial effects of combining phage OMKO1 with anti-pseudomonal antibiotics from five diverse classes. These effects include synergistic killing in four of the five pairings, as well as reduction of antibiotic MIC and suppression of resistant regrowth over 24 hours in all. These findings highlight the flexibility and complexity of phage-antibiotic combinations and call for continued investigation of phage-antibiotic dynamics to optimize synergy. Testing these combinations *in vitro* with clinical samples and evaluating their *in vivo* efficacy will be necessary steps in thoroughly characterizing phage-antibiotic combination treatments and how they may reflect clinical scenarios, as well as ascertaining the depths of their potential to be commonly used adjuvants to antibiotics in a post-antibiotic era.

## Methods

### Experimental Design

PAS assays were conducted using a 96-well plate checkerboard design as illustrated in **Fig. 1**. Each well contains different concentrations of phage and antibiotic. Row A contains the highest concentration of antibiotic which was then serially diluted 2-fold horizontally. Column 1 contains the highest concentration of phage OMKO1. This concentration was serially diluted 1:10 vertically. Column 11 was treated with antibiotic only and Row H was treated with phage only. Column 12 contains treatment and sterility controls. Phage OMKO1 was paired with five antibiotics: Ciprofloxacin, Aztreonam, Tobramycin, Aztreonam, and Colistin. Optical density (OD_600_) values were recorded at 10-minute intervals for 24 hours. Heatmaps represent the average of three biological replicates.

### Bacterial Strain Construction and Maintenance, Growth Media Preparation, and Phage Methodology

*Pseudomonas aeruginosa* strain MPAO1pp-was generated following a *Saccharomyces cerevisiae* recombination technique for construct synthesis and introduced into MPAO1 via conjugation as described in Shanks et al. (67). gDNA was extracted from successful double recombinants following a standard phenol chloroform protocol, prepared using an Illumina Nextera DNA Library Prep Kit, and sequenced by the Yale Center for Genome Analysis (YCGA). Whole genome sequencing confirmed the absence of pf4 and pf6 prophages from MPAO1. Bacterial isolate MPAO1pp-was maintained at -80 °C and revived on Tryptic Soy plates (1.5% agar w/v). Tryptic soy broth (TSB) was prepared according to Sigma Aldrich Millipore (22092-500G) manufacturer protocols with sterilization at 121°C by autoclave for 30 min. TSA was used for all experiments and fresh media was prepared for each replicate. To prepare each overnight culture, a single colony of MPAO1pp-was picked and submerged in 5 mL TSB in a 25 mL flask. Bacteria were incubated at 37°C with agitation at 250 RPM in an Amerex Instruments floor shaker.

Phage OMKO1 was isolated from a freshwater lake sample of Dodge Pond in East Lyme, Connecticut, USA prior to these experiments (63). OMKO1 then underwent whole-genome sequencing analysis and was determined to be a part of family *Myoviridae* genus: *phiKZ-like-viruses* with a size of ∼278 kb. Phage OMKO1 was amplified on *Pseudomonas aeruginosa* strain MPAO1pp-(6). To amplify the phage, 10 µl of a high-titer phage OMKO1 stock was added along with 100 µl of MPAO1pp-overnight culture to 10 mL of TSB. This mixture was left to incubate with agitation (250 RPM) for six hours at 37°C. Following this enrichment, the culture was pelleted by centrifugation (4,000 RCF) and was filtered through a 0.22 µm filter. This phage filtrate was stored at 4°C and was used for each replicate of the experiment.

OMKO1 was titered using the full-plate method. A 1:10 dilution series from undiluted to 10^-11^ of OMKO1 was performed in Gibco Phosphate Buffered Saline (PBS) pH 7.4 without calcium chloride and magnesium chloride. Dilutions were then mixed with a 0.3% semi-solid Tryptic Soy Agar (TSA), 100 µl of overnight culture of MPAO1pp, and plated onto 1.5% solid TSA plates. The plates were incubated, inverted, overnight at 37°C. The titer was determined by quantifying plaque forming units per mL (PFU/mL) at an appropriate dilution.

### Antibiotic preparation

Antibiotic stocks were prepared based on solubility. Ciprofloxacin (solubility: 36 mg/mL), Tobramycin (solubility: 50 mg/mL) and Aztreonam (solubility: 100 mg/mL) were dissolved in sterile deionized water. Ceftazidime (solubility 5 mg/mL) and Colistin (solubility: 10 mg/mL) were dissolved in PBS pH 7.4. All working solutions were filter-sterilized and stored at 4°C for the duration of each experiment.

### Determining Minimum Inhibitory Concentration (MIC) values

The minimum inhibitory concentrations for each antibiotic with *Pseudomonas aeruginosa* strain MPAO1pp-were determined using a standard broth microdilution protocol. Using a 96-well plate, a high concentration of antibiotic was applied to the top row and serially diluted 2-fold vertically. A 1:100 dilution of a MPAO1pp-overnight culture was used to inoculate each well. Plates were then incubated overnight at 37°C with continuous double orbital shaking. Each antibiotic MIC was tested in biological triplicate against positive and negative growth controls. MIC was determined by visual assessment of well turbidity following 24 hours of growth.

### Statistical Analysis

Experiments were performed in biological triplicate. Graphs represent the mean ± standard deviation (bars), except for figure four which shows the mean ± minimum to maximum values (whiskers). Statistical significance when determining differences between treatment types was measured with an ordinary one-way analysis of variance (ANOVA) and Tukey’s multiple comparisons test. Statistical significance of the synergy values compared to zero was measured using a one sample t test and Wilcoxon test with a hypothetical mean value of zero. Significance thresholds were as follows: P > 0.12, not significant (ns); P* < 0.033; P** < 0.002; P*** < 0.001.

## Acknowledgements

We would like to thank all members of the Turner and Koff laboratories for their continued support and insight. We also thank the Cystic Fibrosis Foundation (#006478H223 awarded to ELK) and the Department of Ecology and Evolutionary Biology at Yale University for funding this work.

## Author Contributions

Conceptualization: ELK, BKC, JK, PET

Methodology: ELK, BKC, KEK

Investigation: ELK, KEK

Visualization: ELK

Supervision: BKC, DLM, KEK, PET, JK

Writing–original draft: ELK

Writing–review & editing: ELK, DLM, KEK, BKC, PET, JK

