## Supplemental Table 1 for "Evaluating Synergy of Clinically Utilized Phage OMKO1 with Five Antibiotics from Different Classes against *Pseudomonas aeruginosa*"

Table 1 Average Bliss Synergy Scores (Mean ∆R ± standard deviation) and classifications of five antibiotics at varying phage and antibiotic concentrations. MOI ranges from 10^-7^ PFU/CFU to 10 PFU/CFU. Antibiotic concentrations cover range within each phage-antibiotic interacting zone: ciprofloxacin = 0.016 – 0.063 µg/mL, aztreonam = 0.5– 4 µg/mL, ceftazidime = 0.5 – 4 µg/mL, tobramycin = 0.25 – 1 µg/mL, colistin = 0.25 – 1 µg/mL. Bliss scores calculated from PAS assay AUC data (n=3).

| Antibiotic | Concentration  _(µg/mL)_ | MOI  _(PFU/CFU)_ | Bliss Synergy Score  _(Mean ∆R ± standard deviation)_ | Classification |
| --- | --- | --- | --- | --- |
| Ciprofloxacin | 0.063 | 10 | 0.266 ± 0.040 | **S** |
|  |  | 10^0^ | 0.254 ± 0.004 |  |
|  |  | 10^-1^ | 0.256 ± 0.026 |  |
|  |  | 10^-2^ | 0.290 ± 0.023 |  |
|  |  | 10^-3^ | 0.269 ± 0.045 |  |
|  |  | 10^-4^ | 0.250 ± 0.024 |  |
|  |  | 10^-5^ | 0.225 ± 0.016 |  |
|  |  | 10^-6^ | 0.178 ± 0.024 |  |
|  |  | 10^-7^ | 0.195 ± 0.031 |  |
|  | 0.031 | 10 | 0.126 ± 0.145 | **S** |
|  |  | 10^0^ | 0.086 ± 0.168 |  |
|  |  | 10^-1^ | 0.117 ± 0.160 |  |
|  |  | 10^-2^ | 0.138 ± 0.175 |  |
|  |  | 10^-3^ | 0.086 ± 0.186 |  |
|  |  | 10^-4^ | 0.124 ± 0.138 |  |
|  |  | 10^-5^ | 0.089 ± 0.107 |  |
|  |  | 10^-6^ | 0.049 ± 0.083 |  |
|  |  | 10^-7^ | 0.061 ± 0.095 |  |
|  | 0.016 | 10 | 0.102 ± 0.067 | **S** |
|  |  | 10^0^ | 0.069 ± 0.012 |  |
|  |  | 10^-1^ | 0.062 ± 0.043 |  |
|  |  | 10^-2^ | 0.083 ± 0.021 |  |
|  |  | 10^-3^ | 0.136 ± 0.080 |  |
|  |  | 10^-4^ | 0.061 ± 0.031 |  |
|  |  | 10^-5^ | 0.044 ± 0.033 |  |
|  |  | 10^-6^ | 0.000 ± 0.052 | **A (+)** |
|  |  | 10^-7^ | 0.044 ± 0.028 | **S** |

| Antibiotic | Concentration  _(µg/mL)_ | MOI  _(PFU/CFU)_ | Bliss Synergy Score  _(Mean ∆R ± standard deviation)_ | Classification |
| --- | --- | --- | --- | --- |
|  | 4 | 10 | 0.015 ± 0.047 | **S** |
|  |  | 10^0^ | 0.020 ± 0.062 |  |
|  |  | 10^-1^ | 0.024 ± 0.063 |  |
|  |  | 10^-2^ | 0.018 ± 0.043 |  |
|  |  | 10^-3^ | 0.028 ± 0.059 |  |
|  |  | 10^-4^ | 0.015 ± 0.054 |  |
|  |  | 10^-5^ | 0.004 ± 0.050 |  |
|  |  | 10^-6^ | 0.000 ± 0.055 | **A (+)** |
|  |  | 10^-7^ | -0.016 ± 0.045 | **A (–)** |
| Aztreonam | 2 | 10 | 0.226 ± 0.057 | **S** |
|  |  | 10^0^ | 0.218 ± 0.047 |  |
|  |  | 10^-1^ | 0.230 ± 0.070 |  |
|  |  | 10^-2^ | 0.225 ± 0.051 |  |
|  |  | 10^-3^ | 0.241 ± 0.057 |  |
|  |  | 10^-4^ | 0.200 ± 0.072 |  |
|  |  | 10^-5^ | 0.174 ± 0.035 |  |
|  |  | 10^-6^ | 0.185 ± 0.045 |  |
|  |  | 10^-7^ | 0.165 ± 0.031 |  |
|  | 1 | 10 | 0.157 ± 0.056 | **S** |
|  |  | 10^0^ | 0.129 ± 0.115 |  |
|  |  | 10^-1^ | 0.091 ± 0.024 |  |
|  |  | 10^-2^ | 0.079 ± 0.008 |  |
|  |  | 10^-3^ | 0.160 ± 0.062 |  |
|  |  | 10^-4^ | 0.136 ± 0.059 |  |
|  |  | 10^-5^ | 0.106 ± 0.042 |  |
|  |  | 10^-6^ | 0.150 ± 0.031 |  |
|  |  | 10^-7^ | 0.110 ± 0.065 |  |
|  | 0.5 | 10 | 0.022 ± 0.028 | **S** |
|  |  | 10^0^ | 0.081 ± 0.056 |  |
|  |  | 10^-1^ | 0.107 ± 0.120 |  |
|  |  | 10^-2^ | 0.114 ± 0.033 |  |
|  |  | 10^-3^ | 0.092 ± 0.066 |  |
|  |  | 10^-4^ | 0.031 ± 0.051 |  |
|  |  | 10^-5^ | 0.051 ± 0.035 |  |
|  |  | 10^-6^ | 0.100 ± 0.054 |  |
|  |  | 10^-7^ | 0.081 ± 0.021 |  |

| Antibiotic | Concentration  _(µg/mL)_ | MOI  _(PFU/CFU)_ | Bliss Synergy Score  _(Mean ∆R ± standard deviation)_ | Classification |
| --- | --- | --- | --- | --- |
|  | 4 | 10 | 0.012 ± 0.017 | **S** |
|  |  | 10^0^ | 0.014 ± 0.015 |  |
|  |  | 10^-1^ | 0.017 ± 0.017 |  |
|  |  | 10^-2^ | 0.016 ± 0.018 |  |
|  |  | 10^-3^ | 0.016 ± 0.019 |  |
|  |  | 10^-4^ | 0.014 ± 0.017 |  |
|  |  | 10^-5^ | 0.000 ± 0.010 | **A (+)** |
|  |  | 10^-6^ | -0.026 ± 0.003 | **A (–)** |
|  |  | 10^-7^ | -0.042 ± 0.005 |  |
| Ceftazidime | 2 | 10 | 0.094 ± 0.065 | **S** |
|  |  | 10^0^ | 0.095 ± 0.063 |  |
|  |  | 10^-1^ | 0.097 ± 0.065 |  |
|  |  | 10^-2^ | 0.097 ± 0.067 |  |
|  |  | 10^-3^ | 0.097 ± 0.067 |  |
|  |  | 10^-4^ | 0.096 ± 0.063 |  |
|  |  | 10^-5^ | 0.082 ± 0.049 |  |
|  |  | 10^-6^ | 0.051 ± 0.044 |  |
|  |  | 10^-7^ | 0.022 ± 0.034 |  |
|  | 1 | 10 | 0.123 ± 0.022 | **S** |
|  |  | 10^0^ | 0.127 ± 0.025 |  |
|  |  | 10^-1^ | 0.131 ± 0.028 |  |
|  |  | 10^-2^ | 0.130 ± 0.032 |  |
|  |  | 10^-3^ | 0.104 ± 0.059 |  |
|  |  | 10^-4^ | 0.122 ± 0.036 |  |
|  |  | 10^-5^ | 0.097 ± 0.014 |  |
|  |  | 10^-6^ | 0.056 ± 0.028 |  |
|  |  | 10^-7^ | 0.006 ± 0.020 |  |
|  | 0.5 | 10 | 0.136 ±0.048 | **S** |
|  |  | 10^0^ | 0.107 ±0.032 |  |
|  |  | 10^-1^ | 0.102 ±0.015 |  |
|  |  | 10^-2^ | 0.111 ±0.065 |  |
|  |  | 10^-3^ | 0.130 ±0.028 |  |
|  |  | 10^-4^ | 0.173 ±0.011 |  |
|  |  | 10^-5^ | 0.114 ±0.015 |  |
|  |  | 10^-6^ | 0.093 ±0.033 |  |
|  |  | 10^-7^ | 0.117 ± 0.012 |  |

| Antibiotic | Concentration  _(µg/mL)_ | MOI  _(PFU/CFU)_ | Bliss Synergy Score  _(Mean ∆R ± standard deviation)_ | Classification |
| --- | --- | --- | --- | --- |
|  | 1 | 10 | -0.037 ± 0.068 | **A (–)** |
|  |  | 10^0^ | -0.043 ± 0.066 |  |
|  |  | 10^-1^ | -0.055 ± 0.076 |  |
|  |  | 10^-2^ | -0.033 ±0.070 |  |
|  |  | 10^-3^ | -0.057 ± 0.061 |  |
|  |  | 10^-4^ | -0.067 ± 0.059 |  |
|  |  | 10^-5^ | -0.079 ± 0.053 |  |
|  |  | 10^-6^ | -0.090 ± 0.054 |  |
|  |  | 10^-7^ | -0.096 ± 0.038 |  |
| Tobramycin | 0.5 | 10 | 0.062 ± 0.077 | **S** |
|  |  | 10^0^ | 0.094 ± 0.031 |  |
|  |  | 10^-1^ | 0.063 ± 0.039 |  |
|  |  | 10^-2^ | 0.110 ± 0.045 |  |
|  |  | 10^-3^ | 0.101 ± 0.058 |  |
|  |  | 10^-4^ | 0.075 ± 0.077 |  |
|  |  | 10^-5^ | -0.007 ± 0.061 | **A (–)** |
|  |  | 10^-6^ | -0.034 ± 0.049 |  |
|  |  | 10^-7^ | -0.038 ± 0.023 |  |
|  | 0.25 | 10 | 0.066 ± 0.028 | **S** |
|  |  | 10^0^ | 0.007 ± 0.050 |  |
|  |  | 10^-1^ | -0.016 ± 0.013 | **A (–)** |
|  |  | 10^-2^ | 0.077 ± 0.028 | **S** |
|  |  | 10^-3^ | 0.049 ± 0.036 |  |
|  |  | 10^-4^ | 0.034 ± 0.030 |  |
|  |  | 10^-5^ | 0.041 ± 0.029 |  |
|  |  | 10^-6^ | 0.031 ± 0.069 |  |
|  |  | 10^-7^ | 0.003 ± 0.023 |  |

| Antibiotic | Concentration  _(µg/mL)_ | MOI  _(PFU/CFU)_ | Bliss Synergy Score  _(Mean ∆R ± standard deviation)_ | Classification |
| --- | --- | --- | --- | --- |
|  | 1 | 10 | -0.277 ± 0.066 | **A (–)** |
|  |  | 10^0^ | 0.000 ± 0.056 | **A (+)** |
|  |  | 10^-1^ | 0.004 ± 0.056 | **S** |
|  |  | 10^-2^ | -0.002 ± 0.053 | **A (–)** |
|  |  | 10^-3^ | 0.000 ± 0.056 | **A (+)** |
|  |  | 10^-4^ | 0.001 ± 0.054 | **S** |
|  |  | 10^-5^ | -0.007 ±0.048 | **A (–)** |
|  |  | 10^-6^ | -0.026 ± 0.078 |  |
|  |  | 10^-7^ | -0.011 ± 0.054 |  |
| Colistin | 0.5 | 10 | -0.001 ± 0.127 | **A (–)** |
|  |  | 10^0^ | 0.030 ± 0.011 | **S** |
|  |  | 10^-1^ | 0.123 ± 0.034 |  |
|  |  | 10^-2^ | 0.143 ± 0.104 |  |
|  |  | 10^-3^ | 0.120 ± 0.058 |  |
|  |  | 10^-4^ | 0.154 ± 0.089 |  |
|  |  | 10^-5^ | 0.123 ± 0.032 |  |
|  |  | 10^-6^ | 0.022 ± 0.173 |  |
|  |  | 10^-7^ | 0.085 ± 0.054 |  |
|  | 0.25 | 10 | 0.051 ± 0.012 | **S** |
|  |  | 10^0^ | 0.044 ± 0.048 |  |
|  |  | 10^-1^ | 0.070 ± 0.043 |  |
|  |  | 10^-2^ | 0.039 ± 0.026 |  |
|  |  | 10^-3^ | 0.072 ± 0.033 |  |
|  |  | 10^-4^ | 0.079 ± 0.034 |  |
|  |  | 10^-5^ | 0.072 ± 0.034 |  |
|  |  | 10^-6^ | -0.067 ± 0.225 | **A (–)** |
|  |  | 10^-7^ | 0.010 ± 0.012 | **S** |
